# A comparison study between liquid- and vapor-fed anode zero-gap bioelectrolysis cells

**DOI:** 10.1101/2024.12.21.629895

**Authors:** Nils Rohbohm, Largus T. Angenent

## Abstract

Improving microbial electrosynthesis could be one solution for transitioning towards sustainable chemical production, offering a pathway to convert CO_2_ into valuable commodities from renewable energy sources. Therefore, we examined the performance differences between liquid- and vapor-fed anode zero-gap bioelectrochemical cells for electromethanogenesis, utilizing a membrane electrode assembly to enhance mass and ohmic transport. Focusing on CH_4_ and H_2_ production, we compared two ion-exchange membranes with the liquid-fed anode system and selected the best performing ion-exchange membrane for the vapor-fed anode system. Liquid-fed anode systems did not show significant differences in volumetric CH₄ production rates compared to vapor-fed anode systems, although the latter demonstrated advantages in reducing electrocatalyst degradation and maintaining stable cell voltages. The research underscores the need for further optimization to address performance losses and suggests potential for industrial applications of microbial electrosynthesis, highlighting the importance of catalyst protection.

## Introduction

Renewable sources, such as solar and wind, offer an economically viable alternative to fossil fuels for electricity generation. However, their intermittent nature necessitates extensive energy storage, which is achievable through methods, including hydropower, underground pressure storage, or conversion into chemical energy, such as with batteries. Another approach involves converting electrical energy into chemical energy carriers, such as methane gas (CH_4_) or acetate, through the synergy of electrochemistry and biology.^1^ This dual approach holds the potential to mitigate major greenhouse gas emissions while facilitating the storage of renewable energy.

In general, water electrolysis splits water into H_2_ and O_2_ in the first step. Subsequently, in a second step microbes utilize H_2_ to reduce CO_2_ into CH_4_ or commodity chemicals in a distinct bioprocess, which is known as a gas-fermentation system. The amalgamation of these steps occurs within bioelectrochemical cells, defining the field as microbial electrochemistry.^2^ The specific term for the CO_2_ upgrading at the biocathode is microbial electrosynthesis. While integration possibly could streamline the process economically by eliminating a unit process, it is important to realize that H_2_ and O_2_ are still generated within the bioelectrochemical cell via water electrolysis. Integrated bioelectrochemical cells have promising advantages compared to abiotic electrosynthesis systems, such as CO₂ electrolyzers, which rely on metal catalysts. These benefits are especially important given the challenges abiotic systems face with electrocatalyst selectivity, stability, and material availability.^3^

Geppert and colleagues achieved a breakthrough by implementing the electrochemical design of a redox-flow battery, achieving a current density of 3.5 mA cm^-2^ with approximately 30% energy efficiency in CH production.^4^ Despite a short operating period of only one day, their zero-gap design, where electrodes are closely attached to the membrane, effectively reduced resistance and increased current density from 1.0 to 3.5 mA cm^-2^ during CH_4_ production compared to other studies.^5^ Scaling up the zero-gap design by stacking individual cells represents potential for industrial applications akin to electrolyzers or fuel cells. Rad et al. were the first to improve the current density to 30 mA cm^-2^ with low cell voltages using a hybrid bioelectrochemical system.^6^ However, they protected their cathodic catalysts with a porous transport layer to avoid contact between the catholyte and microbes.

The membrane electrode assembly comprises gas diffusion electrodes and a catalyst-coated membrane used for a zero-gap design, which is readily commercially available. The membrane electrode assembly ensures optimal mass and ohmic transport while reducing cell ohmic losses. For an efficient electrochemical cell, it is imperative to protect the electrocatalysts from contaminants. While abiotic liquid-fed anode electrolyzers necessitate ultra-pure water not to degrade the electrocatalyst, vapor-fed anode electrolyzers only need humid air to resolve a possible source of contamination. This simplifies the overall setup and allows the application of electrolyzers in areas where clean water is not readily available. Using contaminated water or seawater that exhibits low efficiencies with liquid-fed anode electrolyzers would also be possible. Rossi and colleagues enhanced the zero-gap design by introducing vapor into the cathode chamber, using only a phosphate buffer at the anode chamber and utilizing an anion-exchange membrane^7^. Their abiotic system achieved 4.3 mA cm^-2^ in H production without considerably altering the pH gradient between the anode and cathode chambers.

A bioelectrochemical system with a vapor-fed anode was developed to maintain CH₄ production for several weeks at 1.7 mA cm⁻²,^8^ surpassing the operational duration Geppert et al. (2019) reported. The relatively low current density was achieved at a high cell voltage (2.8 V), and further improvements are warranted before commercial viability. Future optimization would undoubtedly improve the current density of their zero-gap design to surpass the current densities compared to configurations with flow electrolyzers or H-cells.

Here, we, firstly, evaluated two different ion-exchange membranes in the liquid-fed anode zero-gap bioelectrochemical cell followed by an investigation of a vapor-fed anode zero-gap cell using the previously selected optimal membrane. Secondly, a comparison was conducted between the liquid-fed and vapor-fed anode systems to evaluate their respective performances. The comparison between the two systems aimed to highlight the benefits of employing a vapor-fed anode, particularly emphasizing its possible potential to prevent pH gradient development and reduce the risk of catalyst wear caused by the liquid flow within the cell. Finally, a vapor-fed anode zero-gap bioelectrochemical cell without and with a PTFE membrane was compared to assess its ability to protect the catalyst layer from degradation caused by exposure to the fermentation broth.

## Results

### Htec-PFSA membrane underperformed compared to the Nafion 117 membrane in terms of volumetric CH_4_ production within liquid-fed anode zero-gap bioelectrochemical cells, thereby, guiding the selection of Nafion 117 for subsequent experimentation in a vapor-fed anode system

Our first aim was to increase the efficiency of bioelectrochemical systems for electromethanogenesis, with a particular focus on optimizing the performance of liquid-fed anode zero-gap cells. We evaluated the effectiveness of two distinct ion-exchange membranes— Htec-PFSA (perfluoro sulfonic acid) and Nafion 117 —within these cells, aiming to ascertain which membrane facilitates the electromethanogenic process.

The experimental procedure involved operating two identical bioelectrochemical cells under liquid-fed anode, zero-gap conditions, with each cell incorporating one of the two selected membranes. These bioelectrochemical systems were operated under a fed-batch mode, receiving a continuous supply of CO_2_ as the only carbon substrate for the electromethanogenesis process. By employing chronopotentiometric control, we ensured a consistent electrical current of 120 mA across the cell, achieving a geometric current density of 7.5 mA cm^-2^ Preliminary findings (not included in this study) had already indicated that such a current density initially produced a cell voltage below 2 V, aligning with one criteria for industrial scalability and efficiency. Each reactor started with an optical density (OD_600_) of 0.04 after inoculation with M. thermautotrophicus ΔH. The inoculation process for the controls followed a sequential approach: first, the reactor with the Htec-PFSA membrane was inoculated, which then served as the inoculum source for the subsequent Nafion 117 membrane reactor.

Upon inoculating the zero-gap bioelectrochemical cell with the Htec-PFSA membrane, a brief one-day lag phase preceded the initiation of electromethanogenesis (Figure 1). Notably, the cell exhibited efficient H_2_ production, achieving a H_2_ Coulombic efficiency (H_2_-CE) of 94.3% (Figure S1). Subsequent volumetric CH_4_ production rate (volume of CH_4_ per bioelectrochemical cell catholyte volume per day), commencing post-lag phase, peaked at 59.9 L L^-1^ d^-1^ by day 10 (Table 1), accompanied by a 74.3% CH_4_ Coulombic efficiency (CH_4_-CE) (Figure 1A and B). Overall, the reactor had a stable phase, displaying minor fluctuations until day 17, maintaining an average volumetric CH_4_ production rate of 47.3 L L^-1^ d^-1^ at a 58.7% CH_4_-CE. The overall average volumetric CH_4_ production rate throughout the 28-day experimental period, excluding the lag phase, was 43.0 L L^-1^ d^-1^ with a 50.8% CH_4_-CE. The maximum energy efficiency (EE) obtained was 36.8% (Figure 1B).

**Figure 1:**
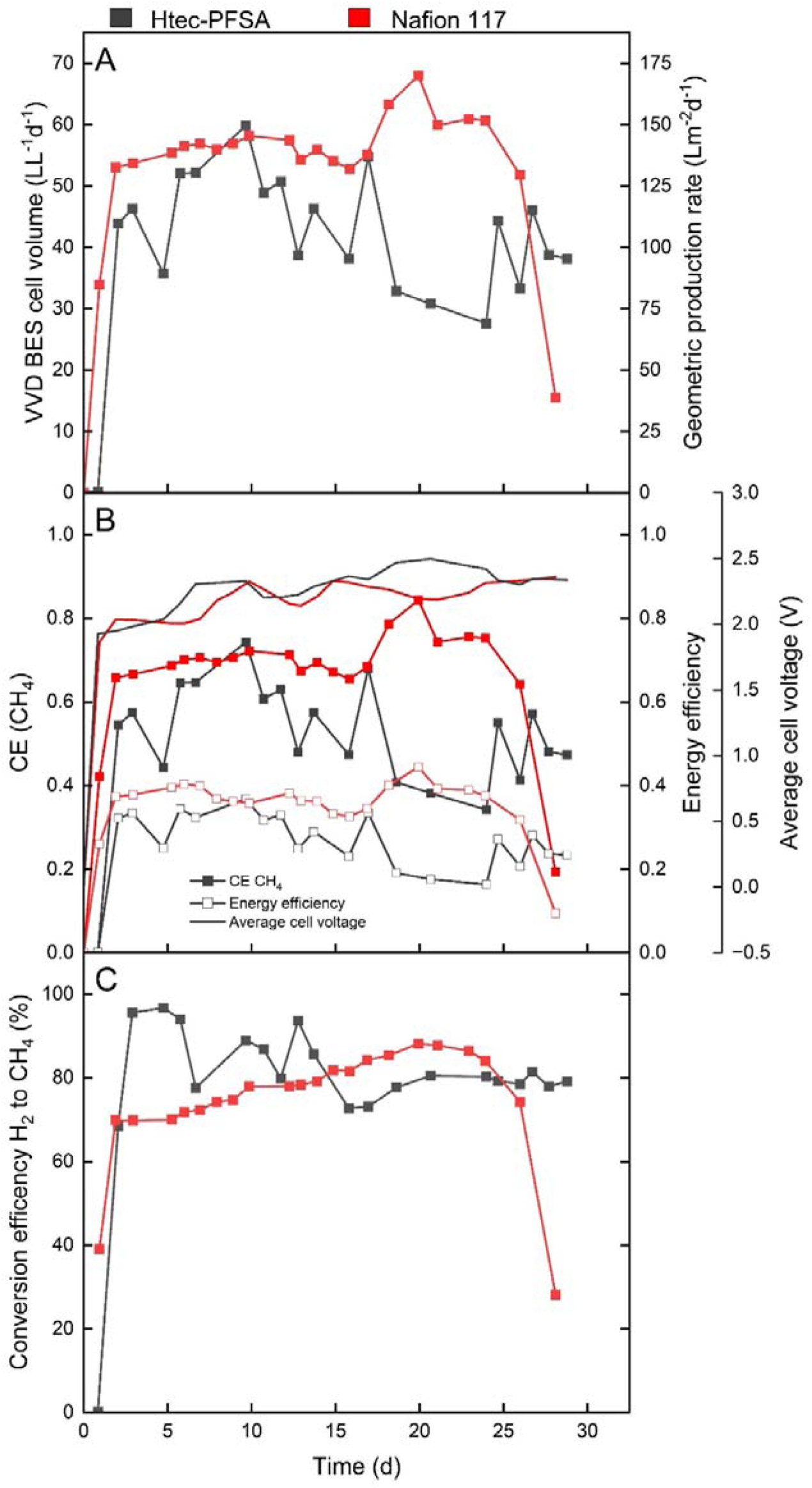
Results from an experiment with a liquid-fed anode zero-gap bioelectrochemical cell using two ion-exchange membranes, Htec-PFSA and Nafion 117, which are coded in dark grey and red, respectively. The different graphs show: (A) the volumetric (Y1-axis) and geometric (Y2-axis) CH_4_ production rate; (B) the CEs (Y1-axis), energy efficiencies (Y2-axis), and average cell voltage (Y3-axis); and (C) the conversion efficiency of H_2_ to CH_4_. The sudden performance drop for the bioelectrochemical cell using Nafion 117 corresponded to a leakage after which we ended the experiment.

**Table 1.** Performance data of the zero-gap bioelectrochemical systems.

| Table 1 Performance data of the zero-gap bioelectrochemical systems. |  |  |  |  |
| --- | --- | --- | --- | --- |
| Zero-gap setups | Absolute current in mA (current density in mA cm<sup>-2</sup>) | Maximum volumetric CH<sub>4</sub> production rate in L·L<sup>-1</sup>·d<sup>-1</sup> | CH<sub>4</sub> Coulombic efficiency in % | Energy efficiency in % |
| Liquid-fed anode with Htec membrane | 120 (7.5) | 59.9 | 74.3 | 36.8 |
| Liquid-fed anode with Nafion 117 membrane | 120 (7.5) | 68.0 | 84.3 | 44.4 |
| Vapor-fed anode with Nafion 117 membrane | 120 (7.5) | 62.5 | 77.5 | 39.7 |
| Vapor-fed anode with Nafion 117 membrane and PTFE membrane at 120 mA | 120 (7.5) | 61.1 | 75.8 | 46.6 |
| Vapor-fed anode with Nafion 117 membrane and PTFE membrane at 400 mA | 400 (25) | 241.8 | 90.0 | 50.2 |

In contrast, the bioelectrochemical cell inoculated with the Nafion 117 membrane experienced no lag phase, likely due to the prior adaptation of microbes to electromethanogenesis. The system demonstrated efficient volumetric CH_4_ production, attaining a peaked volumetric CH_4_ production rate at 68.0 L L^-1^ d^-1^ on day 20 (Table 1), with a CH_4_-CE of 84.4% and a 44.4% EE (Figure 1A and B). The reactor had a stable production phase with minor fluctuations, maintaining an average volumetric CH_4_ production rate of 56.0 L L^-1^ d^-1^ at 69.4% CH_4_-CE, excluding Day 28 when the reactor leaked (drop of volumetric CH_4_ production rate, Figure 1), which also finalized the experiment. Comparing the performance of Htec-PFSA and Nafion 117 ion-exchange membranes in volumetric CH_4_ production reveals an advantage for Nafion 117. Nafion 117 outperformed the Htec-PFSA membrane, showing an 11.9% higher peak volumetric CH_4_ production rate and a 23.1% increase in the average volumetric CH_4_ production rate. This outcome suggests that Nafion 117 has an advantageous application compared to the Htec-PFSA membrane. We hypothesized that the cell exhibited greater stability with the Nafion 117 system, which was further explored in the subsequent section.

### Nafion 117 membrane exhibits better stability and efficiency compared to the Htec-PFSA membrane, evidenced by lower increases in cell voltage and better H_2_ production efficiency

One of the biggest flaws in bioelectrochemistry is the lack of protection of the electrocatalysts, which are susceptible to degradation throughout time due to harsh environmental conditions. These conditions often involve exposure to a medium that is enriched with metals, which are essential for microbial growth, adversely affecting the integrity of the electrocatalysts. While anode degradation can be somewhat mitigated through the utilization of pure water, the inevitable wear and the potential crossover of ions from the cathode to the anode chamber can compromise the performance of anodic electrocatalysts as well. Still, a predominant source of catalyst degradation is attributed to the catholyte, which contains the growth medium for the microbes. Here, the electrocatalyst was Pt/C, which is particularly vulnerable to poisoning by sulfides present in the medium. A key indicator of this degradation is the increase in the cell voltage that is required to sustain a constant applied current, where an increase in cell voltage is inversely related to the activity of the catalyst. Throughout the duration of the liquid-fed anode experiments, both the Htec-PFSA and Nafion 117 membranes exhibited a decline in catalytic activity, which is shown by an increase in cell voltages. Specifically, the Htec-PFSA membrane experiment recorded a cell voltage rise from 1.92 V to 2.49 V (Figure 1B). Conversely, the setup with the Nafion 117 membrane demonstrated a more modest increase, from 1.86 V to 2.36 V. This smaller increase in cell voltage, which we observed with the Nafion 117 membrane, is an advantage for the H_2_ production efficiency.

The ideal scenario involves producing H_2_ without byproducts or losses. However, as cell voltages increase, there is a discernible decrease in catalytic activity through catalyst degradation, increased overpotentials, formation of side products, and loss of selectivity, impacting H_2_ production. The Htec membrane exhibited a higher cell voltage compared to Nafion 117, leading to a steady decline in H_2_-CE and limiting CH_4_ production. Conversely, Nafion 117 demonstrated a more stable H_2_-CE at 91.2% versus 66.5% of the Htec membrane (Figure S1). While both membranes experienced periods of elevated cell voltages, the bioelectrochemical cell with Nafion 117 had an overall lower cell voltage throughout the experiment compared to the cell using the Htec-PFSA membrane, resulting in less disruption to H_2_ production. Notably, at an average cell voltage of 2.47 V, the H_2_-CE of the Htec membrane irreversibly crashed. While the Htec membrane demonstrated a higher conversion efficiency of available H₂ to CH₄ at 82.4% compared to 76.6% of Nafion 117 (Figure 1C), the Nafion 117 setup outperformed overall due to its higher volumetric CH₄ production rate (Table 1) and H₂-CE. As a consequence of the liquid-fed anode zero-gap bioelectrochemical cell performance, the vapor-fed anode experiment was conducted with the membrane electrode assembly using the Nafion 117 ion-exchange membrane.

### Electromethanogenesis in a vapor-fed anode zero-gap bioelectrochemical cell is influenced by pressure adjustments in the anodic chamber, affecting CH_4_ production rates and conversion efficiency

Here, a vapor-fed anode zero-gap bioelectrochemical cell was operated in the same manner as for the liquid-fed anode experiments. The introduction of vapor into the anode chamber was controlled at 3 mL min^-1^, following a 10-min equilibration period, to ensure system stability before commencing measurements. The experiment started with an OD_600_ of 0.04 after inoculation with M. thermautotrophicus ΔH. Similar to the liquid-fed anode experiment utilizing the Htec-PFSA membrane, a one-day lag phase was observed before the onset of electromethanogenesis. During the initial 5 days, without pressurization at the anode, the cell averaged a volumetric CH_4_ production rate of 45.6 L L^-1^ d^-1^, peaking at 49.1 L L^-1^ d^-1^ (Figure 2A). The CH_4_-CE averaged 56.6%, reaching a maximum of 60.9% (Figure 2B). This performance was comparable to that observed with the Htec membrane experiment. To address the issue of back pressure from the cathodic chamber and to reduce H_2_ crossover, the anode chamber was subsequently pressurized to 100 mbar. This adjustment maintained an average volumetric CH_4_ production rate of 47.4 L L d with an average CH_4_-CE of 58.8%. The peak volumetric CH_4_ production rate and CH_4_-CE were 52.2 L L^-1^ d^-1^ and 64.8%, respectively. The highest EE was 39.1% on day 7. More work was required to conclude whether significant differences occurred from pressurization.

**Figure 2:**
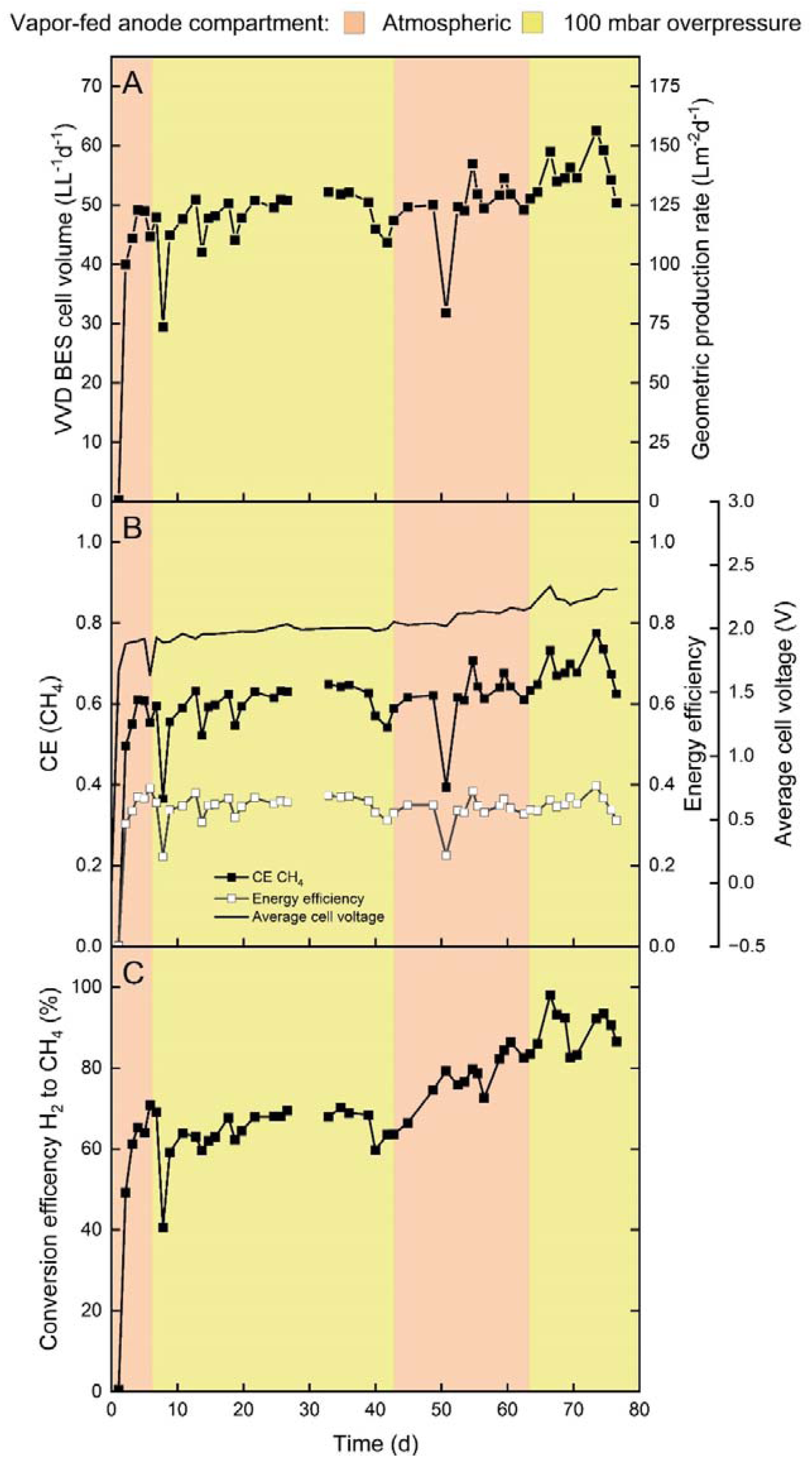
Results from an experiment with a vapor-fed anode zero-gap bioelectrochemical cell using Nafion 117. The anode compartment of the bioelectrochemical cell was kept at atmospheric pressure (orange) or 100 mbar overpressure (yellow). The different graphs show (A) the BES cell volumetric (Y1-axis) and geometric (Y2-axis) CH_4_ production rate; (B) the CEs (Y1-axis), energy efficiencies (Y2-axis), and average cell voltage (Y3-axis); and (C) the conversion efficiency of H_2_ to CH_4_. The decrease in H_2_ production (Figure S2) decreased the excess H_2_ in the system, which resulted in the steady increase in the conversion efficiency of H_2_ to CH_4_.

On day 43, the pressure relief valve was removed to evaluate its impact on system performance. Following removal, the average H_2_-CE was 91.4%, and the H_2_-to-CH_4_ conversion efficiency was 65.5% (Figure 2C, S2), suggesting that H_2_ availability was not a limiting factor, with primary losses attributed to outgassing. Despite the initial high H_2_-CE of 92.7%, it declined to 75.9% by the end of the observation period, averaging 78.7% (Figure S2). Volumetric CH_4_ production remained steady at 49.7 L L⁻¹ d⁻¹. ANNOVA analysis (p>0.05) confirmed that pressurizing the anode chamber to 100 mbar did not improve CH_4_ production rates; however, the H_2_-CE significantly differed (p<0.05) without the pressure relief valve.

On day 51, a sharp drop in CH_4_ production coincided with the highest observed optical density (O.D.) of 0.85 (Figure S3), suggesting nutrient limitations for the microbes. Switching to continuous operation at a hydraulic retention time of 7 days stabilized the O.D. and improved performance. During continuous operation, removing the pressure relief valve increased the average volumetric CH_4_ production rate to 51.6 L L⁻¹ d⁻¹ compared to semi-batch conditions. Reintroducing the pressure relief valve significantly enhanced production (ANNOVA, p<0.05)., averaging 55.7 L L⁻¹ d⁻¹ with a peak of 62.5 L L⁻¹ d⁻¹ and an EE of 39.7% (Table 1).

The H_2_-to-CH_4_ conversion efficiency rose from 80.3% pre-pressurization to 89.8% post-pressurization (Figure 2C), attributed to reduced H_2_ excess in the headspace after day 42. However, H_2_-CE did not significantly change (p>0.05) upon repressurization, and previously high H_2_-CE values were not restored, suggesting lasting system alterations. Continuous operation with a pressurized anode chamber improved both CE and volumetric CH_4_ production rates. However, we cannot rule out that low initial humidity levels, potentially corrected by pressurization, played a pertinent role.

### Vapor-fed anode zero gap bioelectrochemical cell maintains a lower cell voltage and reduces electrocatalyst degradation

In the introductory section, we highlighted the advantages of using a vapor-fed anode bioelectrochemical cell, particularly in circumventing the pH gradient, which contributes to an elevated thermodynamic cell voltage. Additionally, this approach considerably reduces the degradation of the electrocatalyst, which is a notable advantage compared to the liquid-fed anode system, where electrocatalyst degradation is more pronounced. In comparison, the liquid-fed anode bioelectrochemical cell demonstrated a marked increase in cell voltage, from 1.86 V to 2.36 V throughout 28 days (Figure 1B). In contrast, the vapor-fed anode bioelectrochemical cell exhibited voltage stability, maintaining a more stable cell voltage of 1.96 V for 42 days (Figure 2B). This stability persisted until the overpressure was removed from the anode compartment. Subsequent to the adjustment, the cell had a decline in H_2_ production efficiency accompanied by an increase in cell voltage, which peaked at an average maximum of 2.33 V. The stability of the catalyst, which is a factor in the long-term performance of bioelectrochemical systems, was assessed by monitoring the change in cell voltage throughout the operating period and was 172 µV h^-1^.

The degradation of catalysts can change the resistance of the electrolyzer. We evaluated the ohmic resistance involving various contributors such as contact resistance, catalyst resistance, and solution resistance. We employed the current interrupt method to measure the ohmic resistance of the bioelectrochemical cell, as inserting a reference electrode for impedance measurements in the bioelectrochemical cell was not feasible. An increase in bioelectrochemical cell voltage of 500 mV theoretically indicates a rise in ohmic resistance. Our findings corroborate this theory: the initial area-specific ohmic resistance was measured at 18 ± 0.46 Ω cm² at the onset of the experiment and increased to 28 ± 0.10 Ω cm² by its conclusion, which is a 1.5-fold increase. This rise in area-specific ohmic resistance suggests that the membrane electrode assembly has become less efficient. Given that the electrolyte composition remained unchanged (Figure 3). Suggesting minimal impact from solution resistance, we hypothesize that the catalyst and contact resistance are the primary contributors. The liquid-fed anode zero-gap bioelectrochemical cell exhibited an area-specific ohmic resistance of 34 ± 0.33 Ω cm², which was 1.2 times higher at an operation of 28-days versus the 76-days for the vapor-fed anode system. In conclusion, vapor-fed anode bioelectrochemical cells offer advantages in terms of voltage stability and reduced electrocatalyst degradation compared to liquid-fed anode bioelectrochemical systems.

**Figure 3:**
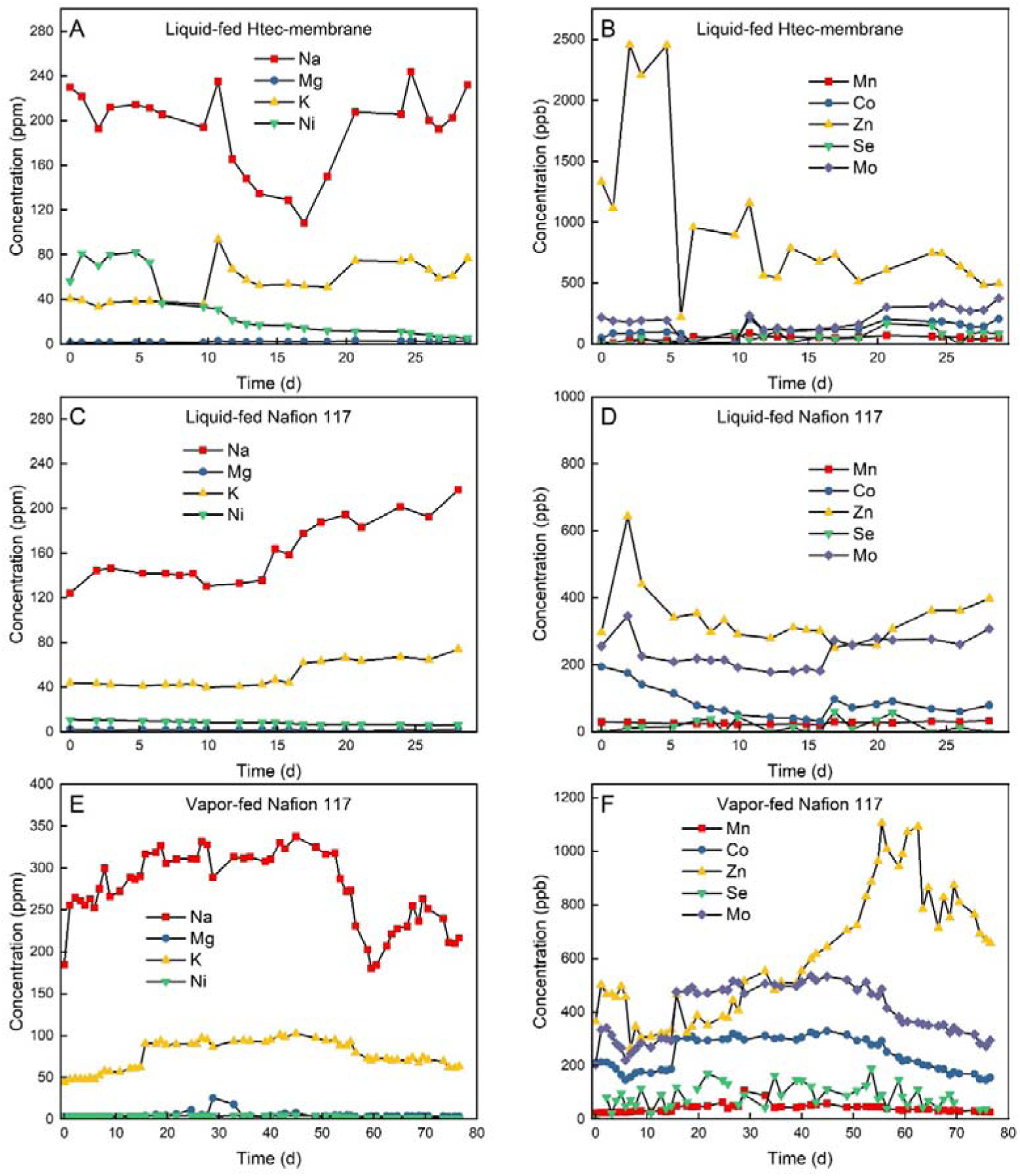
Element concentrations Na, Mg, K, Ni, Mn, Co, Zn, Se, and Mo in the catholyte measured with ICP-MS during the liquid- and vapor-fed anode bioelectrochemical experiments. (A) and (B) shows the liquid-fed experiment using the Htec-PFSA membrane, (C) and (D) demonstrate the liquid-fed anode experiment using the Nafion 117 membrane, and (E) and (F) exhibit the vapor-fed anode experiment using the Nafion 117 membrane.

### Metal compound concentrations did not change during liquid- and vapor-fed anode bioelectrochemical experiments

The zero-gap bioelectrochemical cell experiments observed a consistent decline in H_2_ production, concurrently impeding CH_4_ conversion. This reduction in H_2_ generation indicates reduced electrocatalytic activity at the cathode. Alongside, increasing cell voltage across all experiments further suggests a decline in H_2_ evolution reaction efficacy. The possibility of cathode poisoning via electroplating of metal compounds is considered, which was shown by another study that electroplated metal compounds onto cathodes using concentrated trace element solutions^9^. Consequently, in our study, analytical measurements of various elements, namely ^23^Na, ^24^Mg, ^39^K, ^55^Mn, ^59^Co, ^60^Ni, and ^95^Mo, were conducted (Figure 3). No deficiencies in the medium were noted, and element concentrations remained stable throughout the experiments. Notably, despite no initial deficiency, a decline in ^60^Ni concentration was observed, suggesting an increased uptake by M. thermautotrophicus ΔH compared to other elements. Nickel is used in the Ni-enzymes of M. thermautotrophicus ΔH to enhance growth.^10,11^ While the electroplating of nickel cannot be excluded, the absence of similar trends in other elements challenges this hypothesis. Therefore, the vapor-fed anode bioelectrochemical system is the optimal setup for this study, but measures should be taken to prevent element deficiencies during the operation, especially from nickel and magnesium.

### PTFE membrane, which are placed before the cathode, only gives a small advantage in protecting the catalysts layer

In this experiment, the vapor-fed anode zero-gap bioelectrochemical cell had a PTFE membrane in front of the cathode catalyst. One of the key challenges in bioelectrochemical systems is the degradation of the cathode catalyst due to exposure to the fermentation broth, which can lead to reduced system performance, higher energy consumption, and faster catalyst degradation. By placing a PTFE membrane in front of the cathode, the aim was to create a physical separation between the cathode and the fermentation broth, reducing direct contact, and thus protecting the catalyst from fouling or corrosion similarly to Rad et al.^6^ However, the results demonstrated that the PTFE membrane did not offer a considerable protective advantage, although it did impact other operating parameters.

Initially, the bioelectrochemical cell was operated for 13 days without a PTFE membrane, under conditions identical to the previous experiment (Figure 2). The vapor feeding was increased to ensure that enough moisture was created and avoid pressurizing the system. During the first 13 days, the system stabilized with a CH₄ production rate averaging 53.8 L L⁻¹ d⁻¹, peaking at 63.2 L L⁻¹ d⁻¹ (Figure 4A). The CH₄-CE averaged 66.8%, with a peak of 78.4%, and the EE reached a maximum of 40.8% (Figure 4B). Performance was comparable to earlier experiments, showing slightly higher CH₄ production and EE, although the cell voltage was elevated compared to the previous run. After 13 days, the bioelectrochemical cell was replaced with an identical setup containing a PTFE membrane positioned between the cathode and fermentation broth, leaving a small gap between cathode and PTFE membrane. With the PTFE membrane in place, the system showed a slight reduction in volumetric CH₄ production, averaging 50.4 L L⁻¹ d⁻¹ and peaking at 61.1 L L⁻¹ d⁻¹ (Figure 4A, Table 1). The CH₄-CE decreased to an average of 62.5%, reaching a maximum of 75.8%. However, the key observation was a reduction in cell voltage from 2.18 V (without the PTFE membrane) to 1.91 V (with the PTFE membrane) (Figure 4B). This voltage decrease positively influenced the EE, which peaked at 46.6% in the cell with the PTFE membrane. Compared to the experiment without the PTFE membrane, the cell voltage (average of 1.91 V) is not substantially lower and does not differ considerably. Overall, a small improvement in the cell voltage can be seen but is too small to showcase the benefit of adding a PTFE membrane.

**Figure 4:**
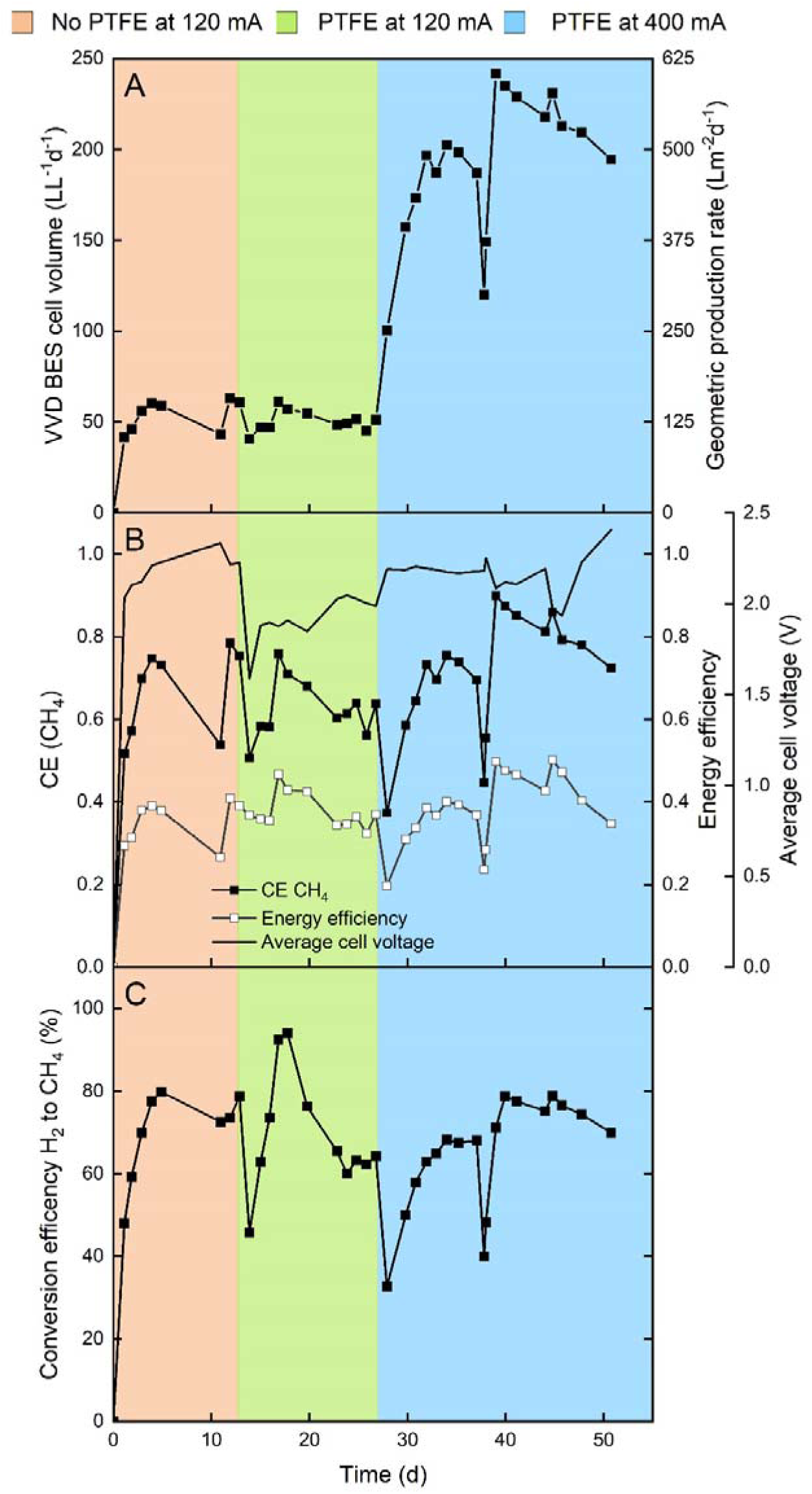
Zero-gap bioelectrochemical vapor-fed anode experiment using Nafion 117. The orange phase represents the initial operation of the bioelectrochemical cell without a PTFE membrane, while the green phase signifies the start of operation with the PTFE membrane added. In the blue phase, the current was increased from 120 mA to 400 mA with the PTFE membrane in place. The different graphs show (A) the BES cell volumetric (Y1-axis) and geometric (Y2-axis) CH_4_ production rate; (B) the CEs (Y1-axis), energy efficiencies (Y2-axis), and average cell voltage (Y3-axis); and (C) the conversion efficiency of H_2_ to CH_4_.

On day 26, the current was increased from 120 mA to 400 mA (7.5 mA cm^-2^ to 25 mA cm^-2^) to assess the response to higher operating stresses. Along with this, the CO₂ feed flow rate was adjusted to 0.8 ml min⁻¹. As a result of the higher H₂ availability, the volumetric CH₄ production rate surged to 241.8 L L⁻¹ d⁻¹, with a CH₄-CE of 90% (Figure 4A and B, Table 1). The average volumetric CH₄ production rate and CH₄-CE were 191.3 L L⁻¹ d⁻¹ and 71.2%, respectively, reflecting an improvement in CH_4_ production efficiency at the higher current. The increase in CH₄-CE compared to the 120 mA operation highlights that raising the current can improve the efficiency in producing CH₄. Interestingly, the fraction of H₂ in the headspace increased at the higher current, and yet the H₂-to-CH₄ conversion efficiency remained similar across both current conditions (Figure 4C). This indicates that the system had more dissolved H₂ available for CH₄ production at the increased current, leading to improved efficiency. This increased efficiency is reflected in the higher energy efficiencies (EEs), with the maximum EE reaching 50.2% (Figure 4B cell voltage 1.97 V, volumetric CH₄ production rate 231.1 L L⁻¹ d⁻¹) on day 44. When the current was increased to 400 mA, the cell voltage initially rose from 1.98 V to 2.18 V. The cell voltage remained stable until day 40, after which fluctuations occurred, with voltages exceeding 3 V being recorded (Figure S4). Such elevated cell voltages can accelerate anode catalyst degradation, making it difficult to maintain long-term voltage stability. However, when the cell voltage was stable, the EE was above 50%, suggesting that the bioelectrochemical cell with a PTFE membrane is capable of maintaining efficient performance and protecting the bioelectrochemical cell from rapid degradation, provided the voltage is well-managed.

### Abiotic tests suggest a crossover of the fermentation broth when protecting the cathode with the PTFE membrane

The goal of incorporating a PTFE membrane in the bioelectrochemical cell was to determine whether it could effectively protect the cathode from degradation while still allowing efficient electron transfer for CH₄ production. Abiotic tests were conducted to isolate the specific impact of the PTFE membrane on the cathode, ensuring that any performance changes were not solely due to biological activity. In these tests, the (bio)electrochemical cell was operated without recirculating the fermentation broth and abiotically, under applied currents of 120 mA and 400 mA, to evaluate how the PTFE membrane impacted the (bio)electrochemical cell performance. Before starting the experiments, vapor was fed to the anode for 1 h. When a current of 120 mA was applied, the cell voltage measured 1.54 V, and at 400 mA, the cell voltage was slightly higher at 1.65 V (Figure S5). The CE-H₂ was 99.9 ± 0.1%, indicating optimal H_2_ production under these conditions.

Comparing the abiotic and biotic experiments revealed considerable differences in cell voltage. At a current of 120 mA, the voltage difference between the abiotic and biotic conditions was 0.37 V, while at 400 mA, the difference increased to 0.53 V. This variation is partly due to the higher thermodynamic potential difference between O_2_ and H_2_ evolution reactions, which is ideally 1.23 V. However, in this setup, the presence of a presumably pH gradient increased the thermodynamic cell voltage during the biotic experiment. Even if the PTFE membrane is hydrophobic, through convection or diffusion, H_2_O from the fermentation broth or from the ion-exchange membrane could have accumulated in the gap between the cathode and the PTFE membrane, which ideally should have contained only H_2_ gas. This accumulation could have created a subsequent pH gradient. However, there is not any substantial proof and given the size of the gap, a collection of a sample was not possible. On day 13 of the vapor-fed anode zero-gap bioelectrochemical cell with the PTFE membrane experiment, the cell voltage was similar to the abiotic experiment at 1.52 V. On day 14, there was a sudden surge in the cell voltage to 1.88 V marking the beginning of the supposed pH gradient. Overall, the PTFE membrane protects the cathode catalyst from the fermentation broth, but due to water convection we assume a pH gradient forming that could be mitigated by increasing the H_2_ production or decreasing the recirculating fermentation broth pressure on the PTFE membrane.

## Discussion

We operated the zero-gap bioelectrochemical experiments successfully with M. thermautotrophicus ΔH under two operating modes, liquid- and vapor-feeding at the anode compartment. While both operating modes have shown comparable volumetric CH_4_ production rates, differences in cell voltages, notably for the vapor-feeding experiment, could make vapor-feeding an option for future studies. The liquid-fed anode experiment had a higher volumetric CH_4_ production rate than the vapor-feeding experiment. Furthermore, the liquid-fed anode experiment reached the highest volumetric CH_4_ production rate and EE, making vapor-feeding the less optimal solution in our study. However, the observations with the liquid-fed anode experiments indicate that degradation is inevitable, and vapor-feeding showed more stable cell voltages and degradation rates. Comparing the volumetric CH_4_ production rate with a similar zero-gap design using vapor-fed anode systems, we had a much higher volumetric CH_4_ production rate while the cell voltage was also lower.^8^ Rad and coworkers managed a higher overall volumetric CH_4_ production rate by protecting their electrocatalyst efficiently at 30 mA cm^-2^ ^6^. In contrast, our study demonstrated a higher EE at 25 mA cm^-2^ by using a PTFE membrane for electrocatalyst protection. To further optimize the zero-gap bioelectrochemical cell, it is necessary to identify the main contributors to electrocatalyst degradation and system performance in both liquid- and vapor-fed anode configurations, providing insights for improved operation in future studies. As an illustration, we had not increased the current density to 25 mA cm^-2^ for the system without the protecting membrane because we anticipated an increase in the cell voltage beyond 2V and a resulting decrease in the EE.

The relatively low H_2_-to-CH_4_ conversion efficiency observed in this study is largely attributed to a higher current density. When low current densities are used, H_2_ is easier miscible in the electrolyte/fermentation broth and can be uptaken by the microbes without forming gas bubbles and outgas.^12^ Geppert and colleagues showed this observation when running a redox flow battery type reactor, which we would classify as a zero-gap cell ^4^. They observed that the outgas concentration of H_2_ increased with higher current densities. However, H_2_ is not an excellent soluble gas, so with higher current densities, more gas bubbles would be formed and are less likely to be uptaken by the microbes. Once the diffusion layer is saturated with dissolved H_2_, H_2_ bubble formation will occur ^12^. Due to the low solubility of H the absolute current needed to saturate the diffusion layer would be reached at low current densities. Here, bubble formation was inevitable because, assuming Fick’s law at 60°C and atmospheric pressure, the minimum current density is 1.37 mA cm^-2^ at which bubble formation would occur.^12,13^ In general, M. thermautotrophicus is an efficient microbe that converts H_2_ and CO_2_ at a very high conversion efficiency and yield, and Electrochaea GmbH now uses it in a commercial setting.^14,15^ Therefore, our study was not limited by the microbe. As for all gas fermentation processes, the low dissolution of H_2_ is the limiting step in efficiently converting H_2_ into any value-added chemical. ^15^ Therefore, our reactor setup presents major flaws in efficiently retaining H_2_ to convert H_2_ and CO_2_ into CH_4_. A solution would be to pressurize the system to dissolve more H_2_ or to recirculate the gas phase to increase the mass transfer rate of dissolved H.^16^ Increasing the surface area of the electrode would enhance the production of miscible H_2_ by providing more active sites for its generation.^17^

PFSA ion-exchange membranes are widely used in electrochemical systems. However, the use of Nafion 117 in this study is not only one of the most studied ion-exchange membranes, it also has a disadvantage. Once the ion-exchange membrane encounters stress, its performance decreases, and H crossover occurs ^18^. Here, the ion-exchange membrane can encounter swelling through exposure to higher operating temperatures, loss of cation-selectivity, mechanical stress from electrolyte recirculation, and ion precipitation on the membrane. While the disadvantages and problems of PFSA ion-exchange membranes are known, it is unclear which disadvantages are the main reason for the performance loss of both liquid- and vapor-fed anode systems. H_2_ crossed over to the anode side (measured qualitatively) in both anodic operating modes, and we hypothesize that the swelling of the membrane is the main contributor to the H_2_ crossover. We recommend testing new ion-exchange membranes that could prevent the above-mentioned disadvantages for future studies.

Furthermore, it has been shown that at higher current densities, more H_2_ crosses through the ion-exchange membrane.^19-21^ The main contributor to H_2_ crossover is the diffusion of supersaturated H_2_. Supersaturated H_2_ is dissolved H_2_ at the boundary layer of the electrode that exceeds the average possible solubility of H_2_, which can be calculated through Henry’s law. Another possibility of H_2_ crossover is convection through electro-osmotic drag or pressure difference. None of all possible H_2_ crossover mechanisms could be shown as a major contributor to both operating modes. While it was impossible to mitigate the H_2_ loss/crossover for the liquid-fed anode bioelectrochemical experiments, we slowed down the H_2_ crossover during the vapor-fed anode experiment by pressurizing the anode compartment to 100 mBar. It was noticeable that the system was stable through the high H_2_-CE and the more or less constant cell voltage until the anodic compartment was set back to atmospheric pressure. However, there is no substantial proof that pressurizing the anodic compartment always improves the performance of the cell. Because pressurizing the anodic compartment improved the humidity levels observed by condensation, the anode might have lacked water for the O_2_ evolution reaction. Higher vapor feeding rates to the PTFE-experiment showed no losses in H_2_ and we observed a H_2_-CE above 90%.

A major contributor to performance loss is the poisoning of the catalyst layer itself. While the Htec-PFSA ion-exchange membrane generally underperformed, the Nafion 117 experiments showed stable H_2_ production in both liquid- and vapor-fed anode operating modes. The difference between the modes is the increased cell voltage, where the vapor-feeding shows its major benefit with its lower and more stable cell voltage during the first 42 days. Several studies have shown that catalyst poisoning is a major contributor to decreased current densities in which the catalyst layer is blocked by precipitate, or the constant electrolyte flow causes abrasion.^6^ The catalyst itself can react with the electrolyte to deactivate its catalytic ability. Here, Pt was inevitably poisoned through sulfide that adsorbs on the active sites of Pt and inhibits the H_2_ evolution reaction.^22^ Other elements that might have been electroplated on the cathode layer showed no concentration reduction except for Ni, which is needed for the methanogen to grow. The other measured elements were protected by a chelating agent (NTA). De Smit and coworkers showed that metals electroplated less on the cathode when a chelating agent was used.^9^ We introduced the PTFE membrane to protect the catalyst, and only small improvements were observed in a lower and more stable cell voltage. It is, therefore, necessary to protect the catalyst layer by separating it better from the electrolyte or using an improved catalyst inert to the electrolyte.

NiMo is stable with methanogen medium while having the same catalytic prowess as Pt.^23,24^ A plate reactor design, utilizing NiMo as cathode catalysts, demonstrated current densities comparable to those observed in this study.^25^ Pentlandite-type catalysts (Fe_3_Co_3_Ni_3_S_8_) also show good catalytic activity and biocompatibility with elevated stability in the presence of H S.^6^ Rad and colleagues, who used pentlandite in their hybrid zero-gap bioelectrochemical cell, showcased high current densities. However, they protected their catalysts with a porous transport layer, which is composed of a PTFE-coated titanium mesh and is commonly used in abiotic electrolyzers to protect the catalysts layer. Similar to their study, we protected the Pt catalyst with a PTFE layer. The performance decreased once the cell voltage increased to higher cell voltages that might have corroded the anode catalyst. In our opinion, the underlying philosophy of bioelectrochemistry is not to protect the catalyst with a hydrophobic layer to prevent contact between the catalyst layer and electrolyte. Instead, the advantage of bioelectrochemistry, compared to the coupling of abiotic electrolysis and gas fermentation, is to have the miscible H_2_ directly taken up by the microbes, rather than releasing H_2_ again as gas in a bubble. Here and elsewehere,^6^ a PTFE layer protected the catalysts. In this case, is microbial electrosynthesis even an option? If high current densities can only be achieved by including a hydrophobic layer between the catalyst layer and electrolyte, an abiotic electrochemical cell can be used instead, and the evolved H_2_ gas can be transported to the bioreactor through a hydrophobic membrane to generate smaller H_2_ bubble size avoiding all the constraints of bioelectrochemical processes. However, a yet unexplored prospect in microbial electrochemistry is using a biofilm at which high current densities without protecting the electrode are theoretically possible.^26^ While creating a biofilm with a high surface area cathode, current densities more than 10 mA cm^-2^ were possible, however, with cell voltages of more than 3 V.^17^

In theory, the benefit of using vapor-fed anode bioelectrochemical cells is the circumvention of a pH gradient that alters the thermodynamic cell potential of water splitting. Ideally, the water vapor at the anode is split into O_2_ and protons. The protons are then transported to the cathode to form H_2,_ leaving the anode compartment at a neutral pH.^27^ Our study showed no improvement in the cell voltage during the vapor-fed anode experiment compared to the liquid-fed anode experiment. Both operating modes had similar starting cell voltages, which diverged to similar final higher cell voltages. The supposedly faster degradation of the electrocatalysts for the liquid-fed anode experiment compared to the vapor-fed anode experiment makes vapor-feeding a better option to protect the catalysts from, for example, abrasion. The abiotic experiment indicated that water accumulation between the cathode and the PTFE membrane likely led to the formation of a pH gradient, resulting in an increase in cell voltage. However, a proof of circumventing a pH gradient by using a vapor-fed anode could not be shown.

## Limitations of the study

The study presents a viable approach for the industrial application of microbial electrosynthesis. However, a considerable challenge remains in the form of low current density and the consequent low EE. Jourdin et al. (2020) postulated that microbial electrosynthesis becomes economically feasible only when the EE exceeds 50%.^28^ Here, the peak EEs recorded were 44.4% for liquid-fed anode and 39.7% for vapor-fed anode experiments (Figure 1B and 2B, Table 1), falling short of the desired threshold. An EE of 50.2% was reached when the cathode was protected with a PTFE layer. A contributing factor to this inefficiency is the excess H_2_ in the headspace, which results in less H_2_ being converted to CH_4_. Recirculating the headspace gas could increase the solubility of H_2_ and increase the EE to more than 50%, making microbial electrosynthesis an option for industrial applications.^16^ Moreover, vapor-fed anode electrolyzers encounter a specific limitation concerning the water content within the membrane.^27^ Low water content in the membrane can lead to increased ohmic resistance. In our experiments, we consistently supplied water vapor to the anodic compartment to maintain high humidity levels inside the cell. However, the humidity levels were not measured, which should be the aim of future studies.

## STAR⍰Methods

### Key resources table

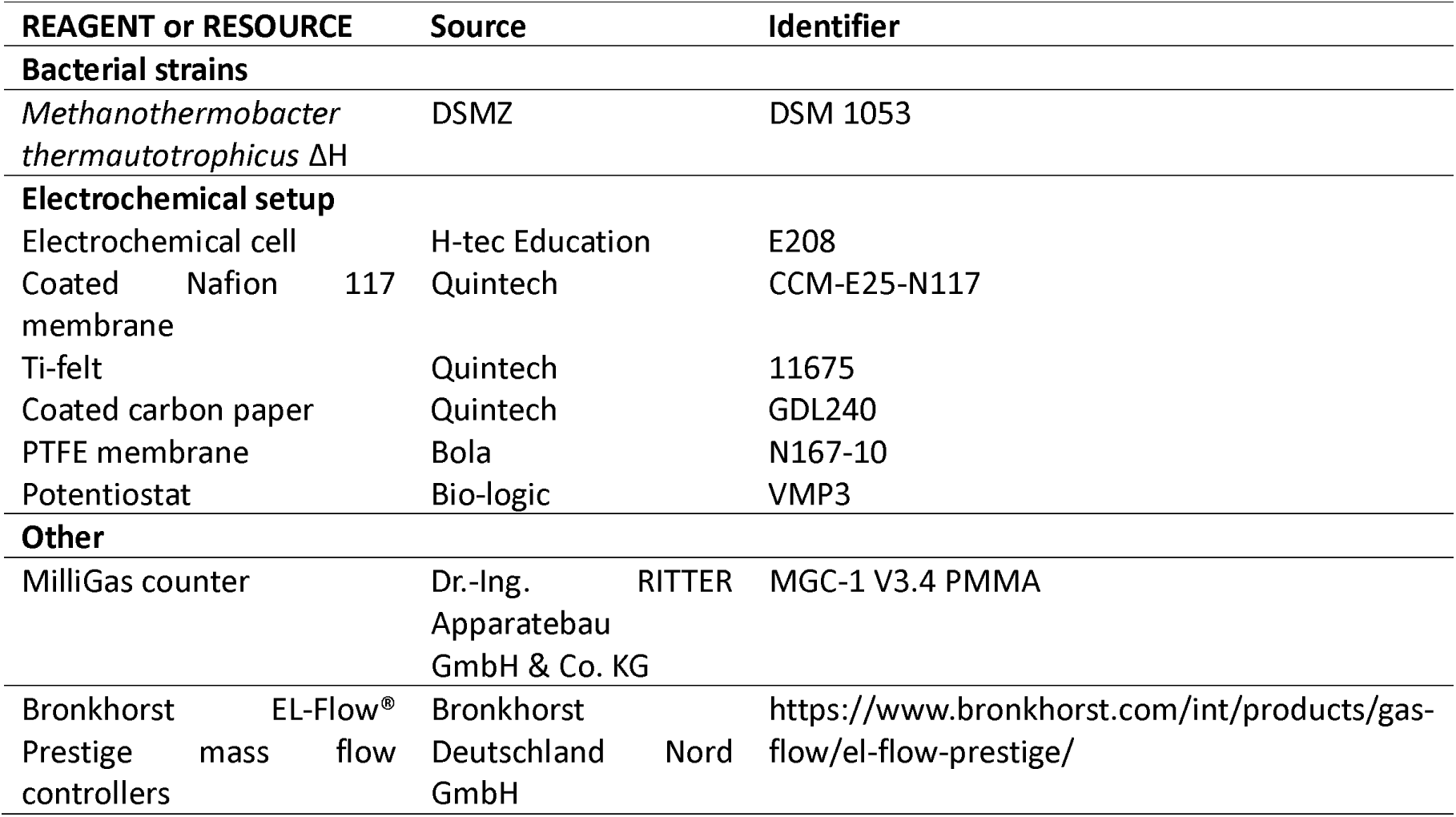

### Resource availability

#### Lead contact

Further information and request for resources should be directed to Largus T. Angenent.

#### Materials availability

This study did not generate new unique materials.

## Experimental Details

### Strain and medium

M. thermautotrophicus ΔH (DSM 1053) was purchased from DSMZ (Braunschweig, Germany). The medium was adjusted from Martin et al.^14^ and contained (per liter): KH_2_ PO_4_, 1.36 g; sodium sulfate, 0.71 g; trisodium nitrilotriacetate, 0.21 g; nitrilotriacetic acid (NTA), 0.076 g; ammonium nickel(II) sulfate, 1.97 mg; cobalt(II) sulfate heptahydrate, 0.71 mg; sodium molybdate dihydrate, 0.61 mg; magnesium sulfate heptahydrate, 0.25 mg; iron(II) sulfate, 0.05 g; ammonium sulfate, 0.06 g; sodium selenate, 0.26 mg; and sodium tungstate dihydrate, 0.32 mg. M. thermautotrophicus ΔH was cultivated for the inoculum in serum bottles, containing 6 g/L sodium bicarbonate as a pH buffer and 0.5 g/L L-cysteine hydrochloride as a reducing agent and sulfur source. The serum bottles were sparged with N_2_/CO_2_ (80/20%, v/v) to create an anoxic environment. Subsequently, the headspace was replaced with H_2_/CO_2_ (80/20%, v/v) at 1 bar overpressure and autoclaved. Incubation of M. thermautotrophicus ΔH occurred at 60°C and 150 rpm. The bioelectrochemical system underwent a similar anoxic treatment by sparging with pure CO_2_ before supplementation and inoculation. Following the establishment of anoxic conditions, the sulfur source (0.3 g/L sodium sulfide) was added, and the pH was adjusted to 7.2 using 1M NaOH, which was diluted in the medium instead of water.

### Bioelectrochemical setup

The electrochemical cell was a commercially available zero-gap electrolysis cell (Figure 5B and C,E208, H-tec Systems GmbH, Augsburg, Germany) composed of a membrane electrode assembly. The perfluorosulfonic acid (PFSA) ion-exchange membrane was coated with 0.5 mg cm^-^² Pt on the cathode side and 2 mg cm^-^² IrRu mix on the anode side. A carbon paper with a mesoporous carbon layer and Ti-felt was placed on top of the catalyst and contacted by the current collector. In subsequent experiments, the frame of the electrolysis cell was kept. However, the catalysts-coated ion-exchange membrane and the gas diffusion layer were exchanged with a Nafion 117 ion-exchange membrane coated with 1 mg cm^-^² Pt/C on the cathode and 2 mg cm^-^² Ir on the anode (CCM-E25-N117, Quintech, Göppingen, Germany). The gas diffusion electrodes were carbon paper with a mesoporous layer and Ti-felt for the cathode and anode side (Quintech, Göppingen, Germany), respectively. A PTFE membrane (N1617-10, Bola, Bohlender GmbH, Grünsfeld, Germany) was used to protect the cathode electrocatalyst. The total geometric surface area of the zero-gap bioelectrochemical cell was 16 cm², and a constant current of 120 mA (7.5 mA cm^-2^) or 400 mA (25 mA cm^-2^) was applied via a potentiostat (VMP3, Bio-logic, Claix, France) and controlled through EC-lab software (Bio-logic, Claix, France). The anode and cathode compartments were connected to glass recirculation vessels (Figure 5A). The catholyte glass recirculation vessel contained multiple ports for pH measurement, gas feed, outgas, in/out ports for the glass recirculation, gas sampling, base feed for pH control, medium feed, and medium out. The anolyte recirculation vessel only contained ports to monitor the pH and recirculate the water to the electrochemical cell.

**Figure 5:**
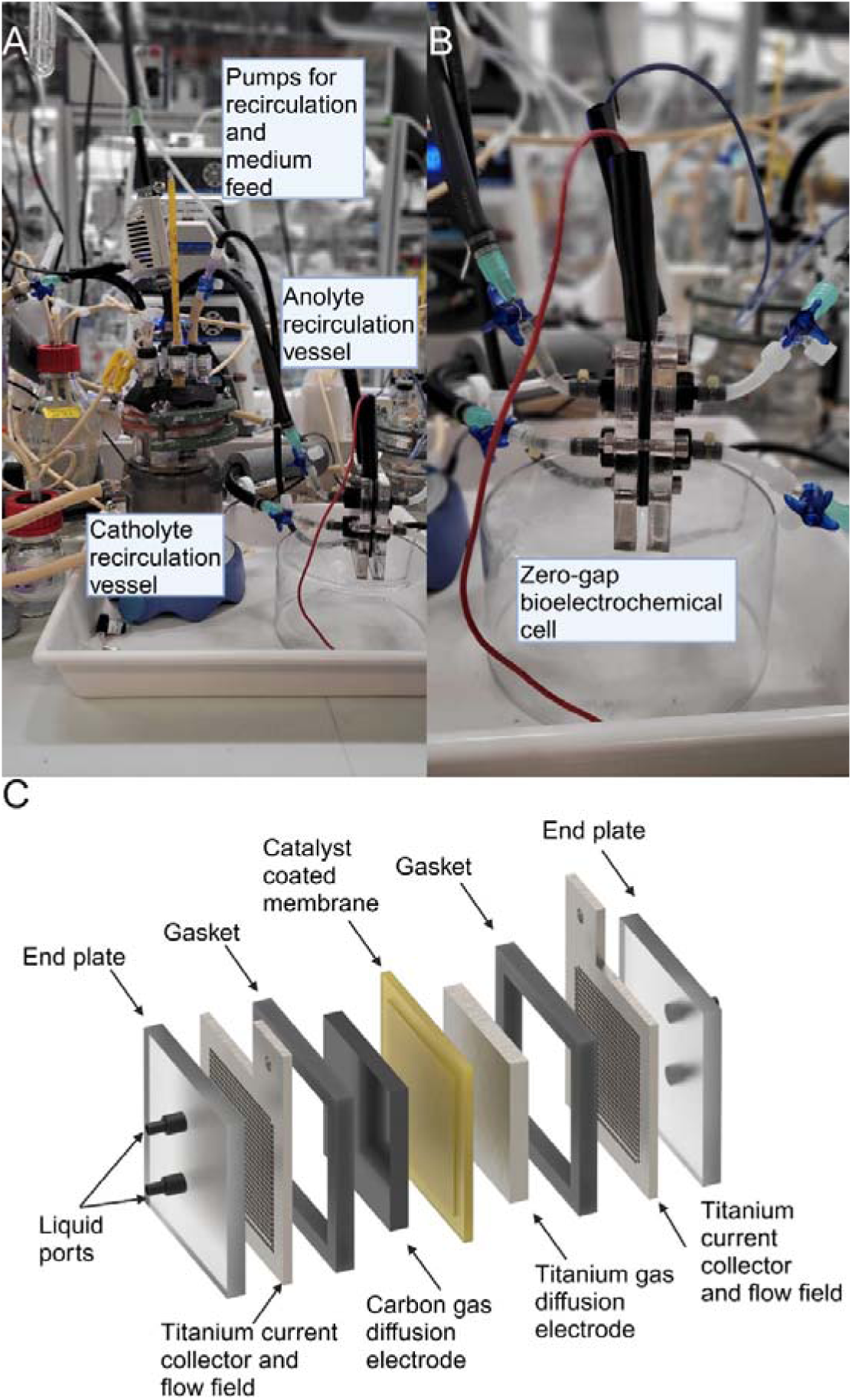
Lab views: (A) Image of the zero-gap bioelectrochemical cell in operation; (B) Close-up view of the zero-gap bioelectrochemical cell; and (C) exploded view of the zero-gap bioelectrochemical cell. The catholyte glass recirculation vessel contained ports to monitor and control the pH and to sample gas. Two ports were used to recirculate the electrolyte/medium. Liquid samples were taken from a three-way valve at the recirculation tubing line. Two ports were used for the gas-in and the gas-out. The gas-out tubing line contained a condenser to avoid water loss. The ports for the anolyte glass recirculation vessel consisted of the recirculation tubing and a pH probe for pH monitoring. During the vapor-fed anode experiment, ports for N_2_ gas-in and a tubing connected to the anodic compartment were used.

During vapor-fed anode operation, water recirculation was replaced by the humid air of the anolyte forced by a N_2_ gas flow rate of 3 mL/min. The anolyte and catholyte were run at 65°C using a water thermostat (KISS 104A, Huber, Raleigh, USA). A multi-channel pump (Masterflex L/S pump equipped with a multi-channel pump head model 7519-20, Cole Parmer, Germany) was used as medium feed in and out during continuous operation (Figure 5A). CO_2_ was used as a carbon source, and the flow rate was controlled using the Bronkhorst EL-Flow® Prestige mass flow controller (Bronkhorst Deutschland NORD GmbH, Kamen, Germany). The CO_2_ flow rate was adjusted from 0.6 mL min^-1^ to 0.3 mL min^-1^ to ensure that carbon is not limited. The outgas volume was measured offline using a MilliGascounter MGC-1 V3.4 PMMA (Dr.-Ing. RITTER Apparatebau GmbH& Co. KG, Bochum, Germany).

Three experiments were conducted with different operating conditions. The first experiment was the commercial electrochemical cell with MilliQ water as an anolyte. The second experiment replaced the ion-exchange membrane with the Pt/Ir-coated Nafion 117 and used MilliQ water as an anolyte. The third experiment was similar to the second experiment, with vapor feeding the anode side. Vapor-feeding was realized by pumping humid air from the anolyte vessel to the anodic side of the bioelectrochemical cell. The cathodic vessel was first sparged with pure CO_2_ for each experiment to make the catholyte/medium anoxic. Before inoculation, the catholyte medium was reduced with 0.3 g L^-1^ sodium sulfide, and the pH was set to 7.2. The inoculation was performed with pre-grown M. thermautotrophicus ΔH in serum bottles to an OD_600_ of approx. 0.04. The bioelectrochemical cells were operated in fed-batch mode. After two weeks, 3 mL of a 100x medium was added. 0.5 mL of 40 g L^-1^ Na_2_S was added every week. The continuous mode operation was exclusively applied to the third experiment and was initiated only after completing the fed-batch phase to compare it with the liquid-fed anode bioelectrochemical cell experiments. The hydraulic retention time (HRT) was 7 days, corresponding to a constant feed rate of 0.03 mL min^-1^. The pH of the catholyte and anolyte was monitored (Bluelab, Gisborne, New Zealand) with a pH electrode (662-1767, VWR, Darmstadt, Germany). A sample day consisted of measuring OD_600_, pH, and out gas composition. The liquid sample was stored at -20 °C until further analysis. The ohmic resistance of the zero-gap bioelectrochemical cell was measured using the current interrupt method using the potentiostat (Bio-logic, Claix, France). A current of 120 mA or 400 mA, which was the applied current during the zero-gap bioelectrochemical experiments, was used during the current interrupt with 1 s intervals repeated 10 times.

## Analytical analysis

A gas chromatograph (SRI GC, Torrance, USA) was used to analyze the headspace gas composition. The gas chromatograph was equipped with a Haysep D column (length 3m, outer diameter 1/8”, SRI GC). A thermal coupled detector and a flame ionization detector were used to measure H_2_, CH_4_, and CO_2_ with N_2_ as the carrier gas. The column temperatures and pressures were set at 70°C and 20 psi, respectively. 500 µL was taken from the gas headspace with a gastight syringe (Hamilton, Reno, USA) for each sample point. The OD_600_ was measured with a UV/VIS-Spectrophotometer at 600 nm absorbance (BioMate™ 160, ThermoFischer Scientific, Waltham, USA). All liquid samples were analyzed on an Agilent 7900 ion-coupled plasma mass spectrometer instrument (Agilent, Santa Clara, USA) to quantify ^23^Na, ^24^Mg, ^39^K, ^44^Ca, ^55^Mn, ^56^Fe, ^59^Co, ^60^Ni, ^66^Zn, ^78^Se, and ^95^Mo. The samples were diluted 1:100 in 1% HNO_3,_ which was also the matrix.

## Calculations

The Coulombic efficiency was calculated by dividing the produced CH_4_ or H_2_ in number of electrons by the number of electrons supplied to the electrochemical cell. The equation is as follows:

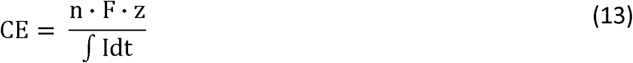

Where n is the molar quantity of produced CH_4_ or H_2_, F the Faraday constant, z number of electrons to form CH_4_ or H_2_, and I the current of the bioelectrochemical cell (7.5 mA cm^-2^). The energy efficiency was calculated as follows:^29^

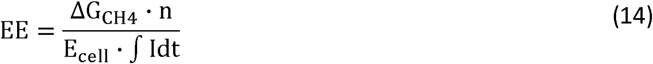

With ΔG_CH4_ the Gibbs free energy of CH_4_ (890.4 kJ/mol), and E_cell_ the measured cell voltage. The conversion efficiency of H_2_ to CH_4_ was calculated as follows:

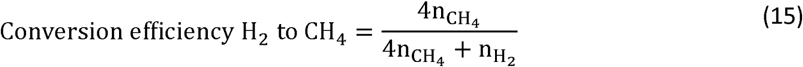

With n_CH4_ the quantity of mole of CH_4_ and n_H2_ the quantity of mole of H_2_.

## Acknowledgments

This work was supported by the Deutsche Forschungsgemeinschaft (DFG, German Research Foundation, EBiotech, SPP2240; L.T.A.) – project number 445506379, the Alexander von Humboldt Foundation in the framework of the Alexander von Humboldt Professorship (L.T.A.), and The Novo Nordisk Foundation CO_2_ Research Center with grant number NNF21SA0072700 (L.T.A.).

## Author contributions

Nils Rohbohm: Methodology, Formal Analysis, Investigation, Data Curation, Writing – Original Draft & Review & Editing, Visualization, Funding Acquisition; Largus T. Angenent: Conceptualization, Resources, Writing –Review and Editing, Supervision, Project administration, Funding acquisition.

