## Supplementary Information for "A comparison study between liquid- and vapor-fed anode zero-gap bioelectrolysis cells"

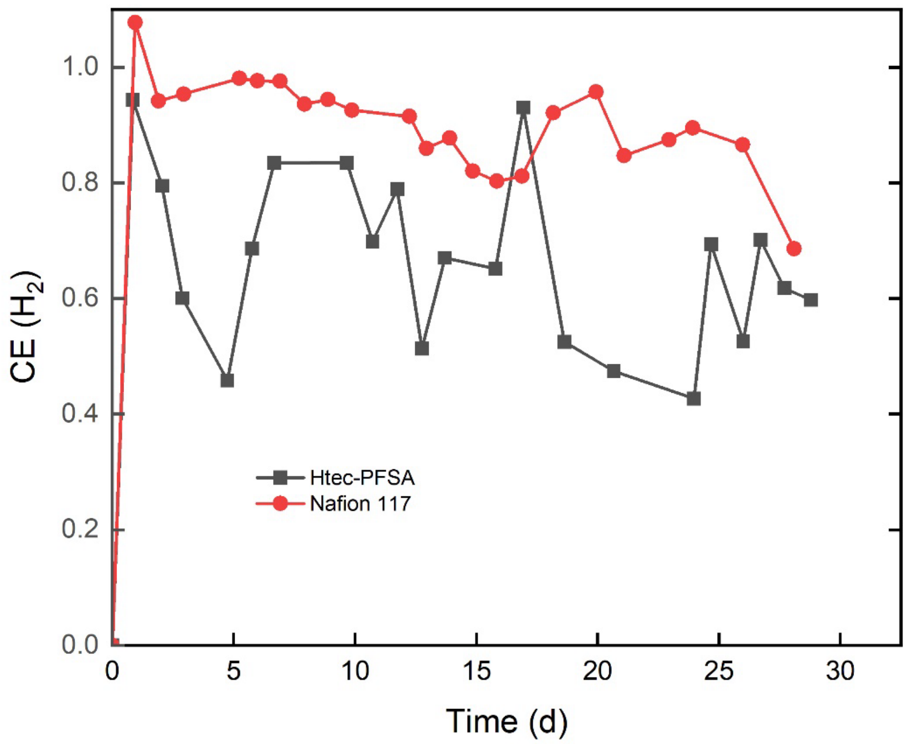


**Figure S1**: The CE of $H_2$ during the liquid-fed anode zero-gap bioelectrochemical cell experiment using Htec-PFSA (dark grey) and Nafion 117 (red) as ion-exchange membrane.

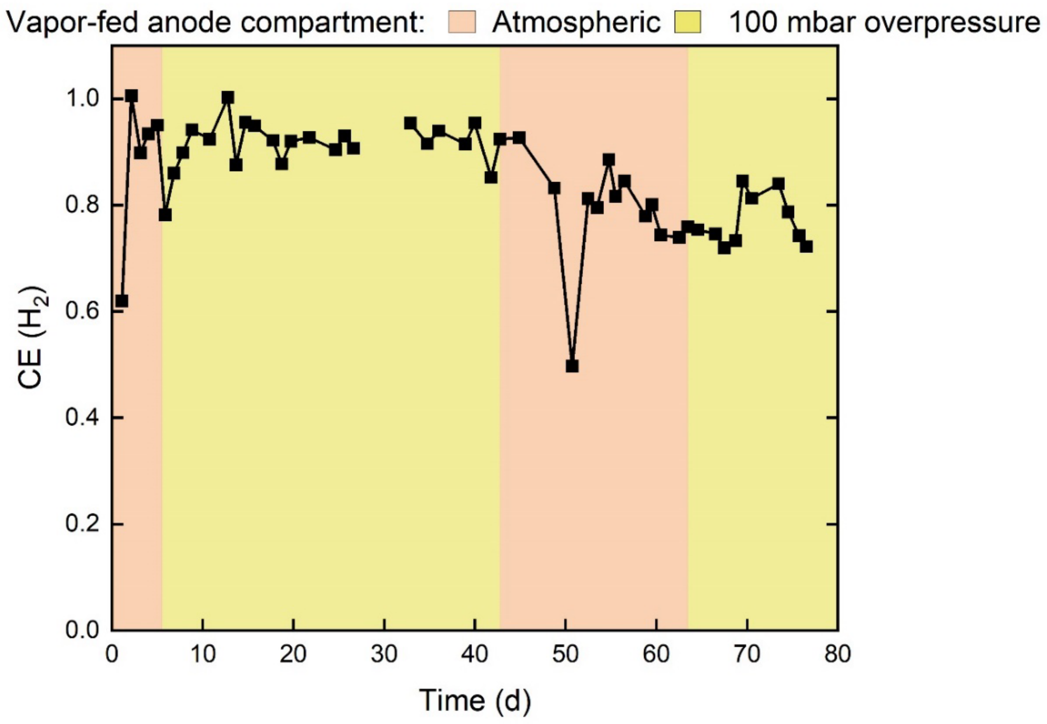


**Figure S2**: The CE of $H_2$ during the vapor-fed anode zero-gap bioelectrochemical cell experiment using Nafion 117 as the ion-exchange membrane. The anode compartment of the bioelectrochemical cell was kept at atmospheric pressure (orange) or 100 mbar overpressure (yellow).

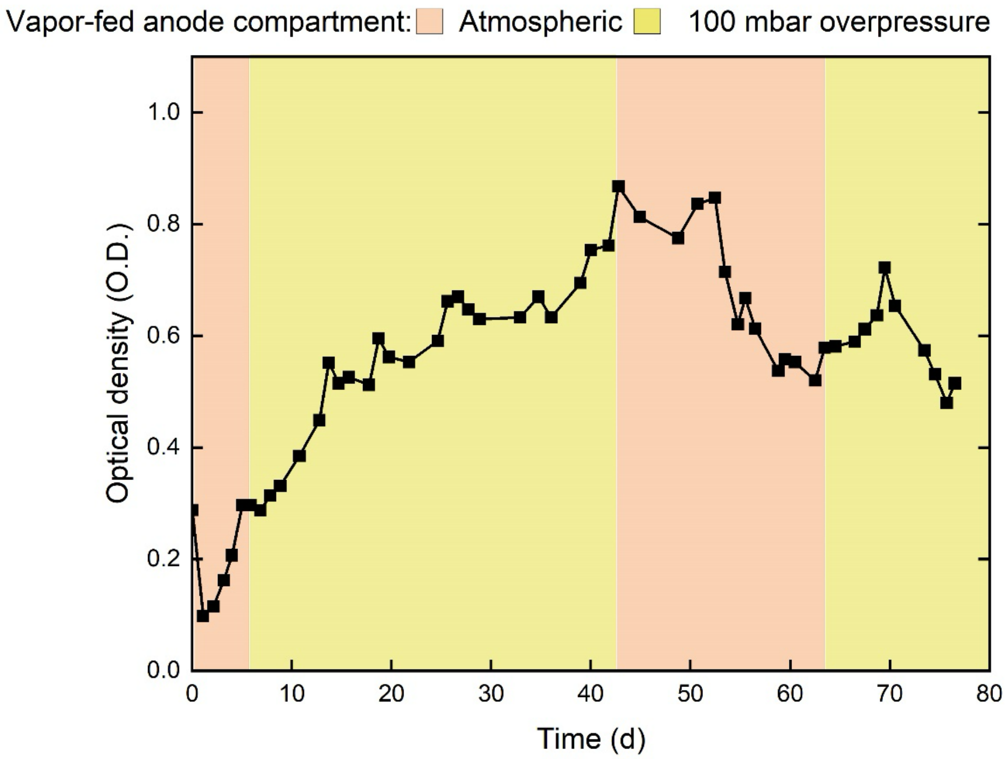


**Figure S3**: The optical density (O.D.) during the vapor-fed anode zero-gap bioelectrochemical cell experiment using Nafion 117 as the ion-exchange membrane. The anode compartment of the bioelectrochemical cell was kept at atmospheric pressure (orange) or 100 mbar overpressure (yellow).

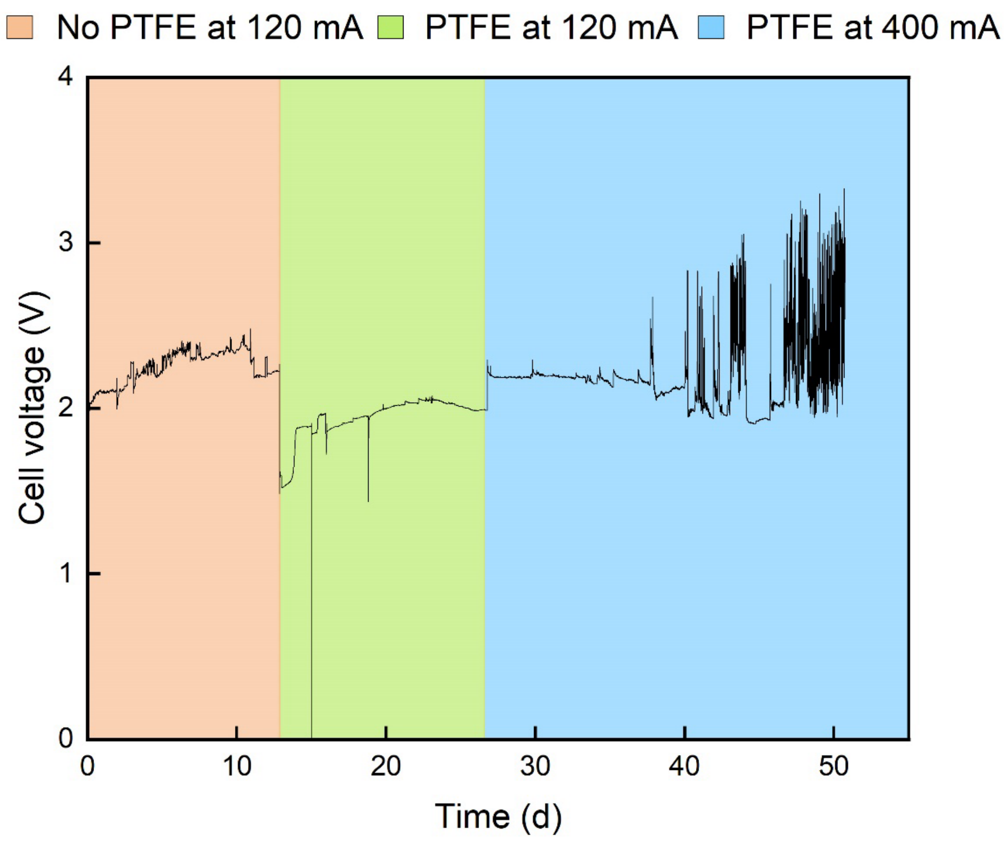


Figure S4: The cell voltage of the vapor-fed anode bioelectrochemical cell, which was run abiotically at 120 mA (green) and 400 mA (blue). The orange phase represents the initial operation of the bioelectrochemical cell without a PTFE membrane, while the green phase signifies the start of operation with the PTFE membrane added. In the blue phase, the current was increased from 120 mA to 400 mA with the PTFE membrane in place.

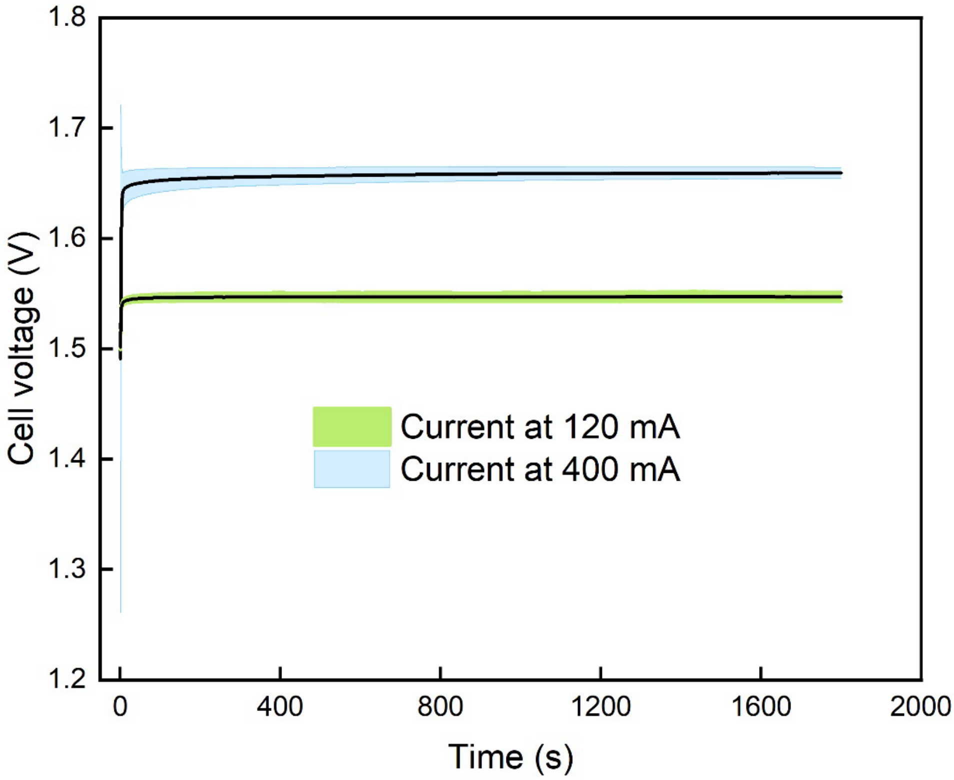


Figure S5: The cell voltage of the vapor-fed anode bioelectrochemical cell, which was run abiotically at 120 mA (green) and 400 mA (blue).
